# Single-cell transcriptomics defines heterogeneity of epicardial cells and fibroblasts within the infarcted heart

**DOI:** 10.1101/2021.01.26.428270

**Authors:** Julia Hesse, Christoph Owenier, Tobias Lautwein, Ria Zalfen, Jonas F. Weber, Zhaoping Ding, Christina Alter, Alexander Lang, Maria Grandoch, Norbert Gerdes, Jens W. Fischer, Gunnar W. Klau, Christoph Dieterich, Karl Köhrer, Jürgen Schrader

**Author notes:** Corresponding author.; Correspondence: Jürgen Schrader, MD, Department of Molecular Cardiology, Heinrich Heine University Düsseldorf, Universitätsstr. 1, 40225 Düsseldorf. These authors contributed equally to this work (Julia Hesse, Christoph Owenier).

## Abstract

In the adult heart, the epicardium becomes activated after injury, contributing to cardiac healing by secretion of paracrine factors. Here we analyzed by single-cell RNA sequencing combined with RNA in situ hybridization and lineage tracing of WT1^+^ cells the cellular composition, location, and hierarchy of epicardial stromal cells (EpiSC) in comparison to activated myocardial fibroblasts/stromal cells in infarcted mouse hearts. We identified 11 transcriptionally distinct EpiSC populations, that can be classified in three groups each containing a cluster of proliferating cells. Two groups expressed cardiac specification makers and sarcomeric proteins suggestive of cardiomyogenic potential. Transcripts of HIF-1α and HIF-responsive genes were enriched in EpiSC consistent with an epicardial hypoxic niche. Expression of paracrine factors was not limited to WT1^+^ cells but was a general feature of activated cardiac stromal cells. Our findings provide the cellular framework by which myocardial ischemia may trigger in EpiSC the formation of cardioprotective/regenerative responses.

## Introduction

Myocardial infarction (MI), still the most frequent cause of death in western societies, is associated with massive activation of cardiac fibroblasts, ultimately resulting in excessive accumulation of extracellular matrix (ECM) components that finally impair cardiac function (1). During development, the majority of cardiac fibroblasts are derived from the epicardium which forms the thin outermost epithelial layer of all vertebrate hearts and exhibits extensive developmental plasticity (2). A subset of epicardial cells undergoes epithelial to mesenchymal transition (EMT) and those epicardial progenitor cells can give rise to various cardiac cell types. In addition to cardiac fibroblasts, this includes vascular smooth muscle cells and pericytes which contribute to the coronary vasculature (2). Embryonic epicardial heterogeneity was recently studied at the single-cell level in the developing zebrafish heart und uncovered three epicardial populations, functionally related to cell adhesion, migration and chemotaxis (3).

In the adult heart, the epicardium is a rather quiescent monolayer that becomes activated after MI by upregulating embryonic epicardial genes (4,5). In the injured heart, epicardial cells form via EMT a multi-cell layer of epicardial stromal cells (EpiSC) at the heart surface that can reach a thickness of about 50-70 µm in mice (6). It is generally assumed that the activated epicardium recapitulates the embryonic program in generating mesenchymal progenitor cells, although there may be major molecular differences with respect to their embryonic counterpart (7). On the functional side, adult EpiSC secrete paracrine factors that stimulate cardiomyocyte growth and angiogenesis (8) and play a key role in post-MI adaptive immune regulation (9). When stimulated with thymosin β4, Wilms tumor protein 1-positive (WT1^+^) adult EpiSC can form cardiomyocytes, however, the rate of conversion is only small (10). Thus, the epicardium is a signaling center regulating cardiac wound healing and may have cardiogenic potential in the adult injured heart.

Despite its importance in cardiac repair, little is known on cell heterogeneity and molecular identifiers within the epicardial layer of the adult heart. Yet, this knowledge is essential to map the epicardial progeny and attribute meaningful functions. While the single-cell landscape of activated cardiac fibroblasts (activated cardiac stromal cells, aCSC) in the post-MI heart has been explored in detail (11,12), these studies did not assess epicardial heterogeneity because of lack of specific identifiers. In a previous study we have reported a novel perfusion-based technique (13) which permitted the simultaneous isolation of EpiSC and aCSC with high yield and only minimal cell activation. The isolation of viable, purified preparations of aCSC and EpiSC from the infarcted heart permitted the first direct comparison of the two cardiac stromal cell fractions. We have combined single-cell RNA sequencing (scRNAseq) with lineage tracing of WT1-expressing cells and localization of cell populations by RNA *in situ* hybridization, characterized cellular hierarchy of adult EpiSC, defined similarities and differences to cardiac fibroblast, and explored in detail the individual EpiSC populations in the activated epicardium.

## Results

### ScRNAseq of post-MI stromal cells

EpiSC (epicardial stromal cells) and aCSC (activated cardiac stromal cells) were isolated from the same mouse hearts (n=3) 5 days after MI (50 min ischemia followed by reperfusion) using a technique which rather selectively removed EpiSC by applying gentle shear force to the cardiac surface (13) (Figure 1A). CSC (cardiac stromal cells) were isolated from uninjured hearts of sham-operated animals (n=3). Cell preparations were depleted of cardiomyocytes, endothelial cells and immune cells (see methods) prior to scRNAseq using the 10x Genomics Chromium platform. Transcriptional profiles of 13,796 EpiSC, 24,470 aCSC and 24,781 CSC were captured after quality control filtering. Unbiased clustering using the Seurat R package with visualization in UMAP dimension reduction plots was performed to identify cells with distinct lineage identities and transcriptional profiles. Average and significantly enriched RNA expression of EpiSC, aCSC and CSC are listed in Supplementary files 1 and 2, respectively.

**Figure 1.**
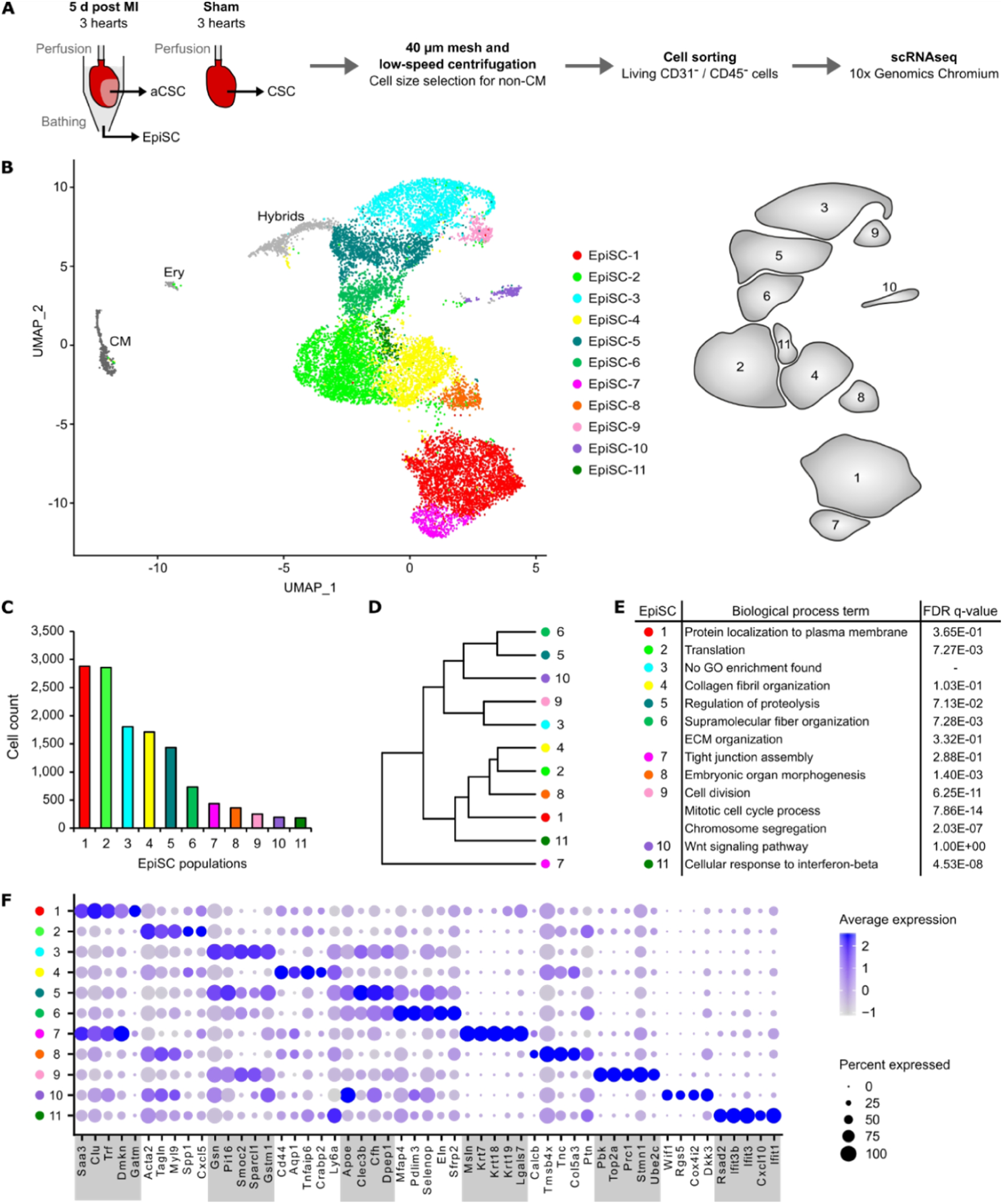
Cell populations in EpiSC from the infarcted heart. (**A**) Schematic workflow. EpiSC and aCSC were simultaneously collected from the surface and the myocardium of the isolated perfused heart by applying mild shear forces to the cardiac surface (13) at 5 day post MI (n=3). CSC were purified from 3 non-infarcted control hearts 5 days after sham surgery. Mesh purification, low-speed centrifugation and cell sorting by flow cytometry was performed to remove cardiomyocytes, CD31^+^ endothelial cells, CD45^+^ immune cells and apoptotic or necrotic cells before analysis using the 10x Genomics Chromium platform. (**B**) UMAP plot of clustered scRNAseq data of the pooled EpiSC fraction (*n*=13,796 single cells). Identified EpiSC populations are color-coded as well as shown in the scheme on the right. CM, cardiomyocytes; Ery, erythrocytes. (**C**) Cell count of EpiSC populations. (**D**) Dendrogram of EpiSC populations according to average RNA expression. (**E**) Significant GO biological process terms. (**F**) Dot plot of top 5 marker genes for each EpiSC population.

### Characterization of EpiSC populations

As shown in Figure 1B, we identified 11 transcriptionally different cell populations within the EpiSC fraction. Cell doublets with hybrid transcriptomes identified by DoubletFinder (16) (Figure 1—figure supplement 1A) and minor non-stromal cell populations such as cardiomyocytes and erythrocytes, identified according to cell type-specific marker expression (Figure 1—figure supplement 1B), were excluded from further analysis. EpiSC populations varied in size (Figure 1C), were hierarchically structured (Figure 1D) and showed over-representations of distinct GO biological process terms (Figure 1E). The top 5 most differentially expressed genes within each population are displayed in Figure 1F. Remarkably, transcripts of epithelial cell-associated genes and established epicardial genes (23) were enriched in both EpiSC-1 (*Dmkn, Saa3*) and EpiSC-7 (*Msln, Krt7, Krt18, Krt19, Lgals7*), while mesenchymal marker genes were primarily expressed in EpiSC-3 (*Gsn, Pi16*) and EpiSC-4 (*Cd44, Ly6a*). Genes coding for ECM proteins were highly expressed in EpiSC-2 (*Spp1*), EpiSC-3 (*Smoc2, Sparcl1*), EpiSC-5 (*Clec3b*), EpiSC-6 (*Mfap4, Eln*) and EpiSC-8 (*Col5a3, Tnc*). Genes encoding contractile proteins were preferentially expressed in EpiSC-2 (*Acta2, Myl9, Tagln*). Wnt pathway-associated gene transcripts were enriched in EpiSC-6 (*Sfrp2*) and EpiSC-10 (*Wif1, Dkk3*). Genes related to the cellular response to interferon characterized EpiSC-11 (*Ifit1, Ifit3* and *Ifit3b, Cxcl10*). Finally, expression of genes associated with high cell cycle activity and mitosis was a feature of EpiSC-9 (*Pbk, Top2a, Prc1, Stmn1, Ube2c*).

### Epicardial marker gene expression

The cellular distribution of well-established epicardial progenitor marker genes such as *Wt1, Tbx18, Sema3d, Aldh1a2, Gata5* and *Tcf21* (2) is shown in Figure 2A. *Wt1*, commonly used for lineage tracing studies (4), was found to be highly enriched in EpiSC populations 1, 7 and 8, which are adjacent to each other (Figure 1B). *Tbx18*, on the other hand, was broadly distributed in populations 1, 2, 4, 7, 8, 9 and 11 (Figure 2A). *Sema3d* was predominantly expressed in EpiSC-7, while *Aldh1a2* mainly resides in EpiSC populations 1, 3, 7 and 9. *Tcf21* again was broadly expressed showing no overlap with *Wt1*-expressing clusters.

**Figure 2.**
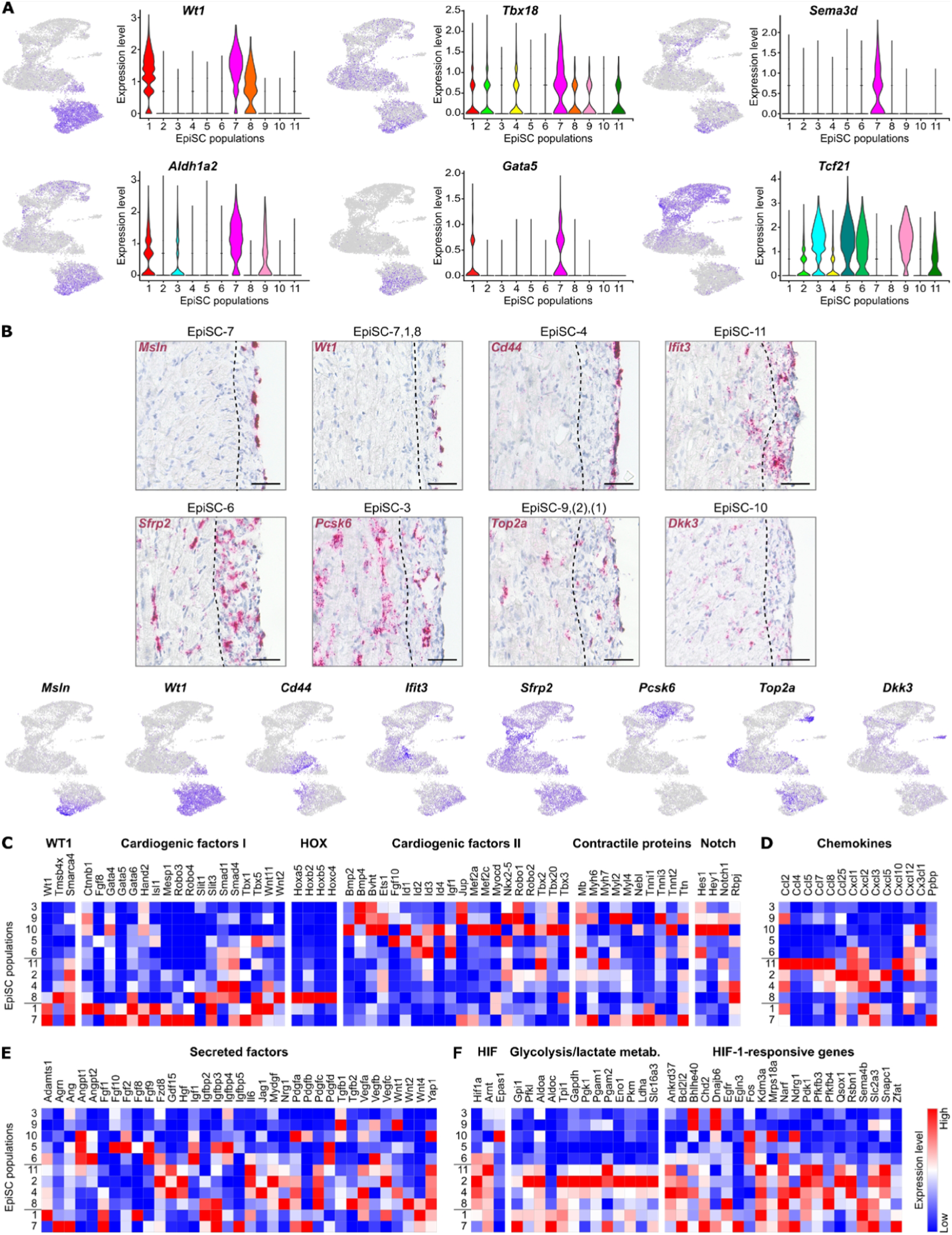
Molecular characterization and location of EpiSC populations. (**A**) Expression of epicardial progenitor cell markers visualized in feature and violin plots. (**B**) Upper panel: RNA *in situ* hybridization of EpiSC population identifiers (red) in heart cryosections 5 days post MI. Representative images (*n*=4 hearts) of the infarct border zone are shown. The dotted line marks the interface between myocardial / epicardial tissue according to cell morphology. Nuclei were stained with hematoxylin (blue). Scale bars, 50 µm. Lower panel: Feature plots visualizing EpiSC population molecular identifiers. (**C** to **F**) Heat maps showing the expression of cardiogenic factors and Notch target genes (C), chemokines (D), further secreted factors (E) as well as genes associated with HIF-1 signaling (F). EpiSC populations are listed according to their position on the UMAP plot.

Since cells in EpiSC-1 showed substantial inhomogeneity of markers (Figure 2A), we carried out a separate clustering analysis for EpiSC-1 including the adjacent two *Wt1*-expressing populations EpiSC-7 and 8. We identified 5 subclusters in EpiSC-1 (EpiSC-1.1, 1.2, 1.3, 1.4, 1.5), with no subclustering in EpiSC-7 and 8 (Figure 2—figure supplement 1A-C). EpiSC1.4 is characterized by expression of cell cycle-associated genes (*Rrm2, Pclaf, Hist1h2ap, Hmgb2, Ube2c, Top2a*), indicating proliferating cells (Figure 2—figure supplement 1D; Supplementary files 3 and 4). EpiSC1.5 showed enriched expression of genes encoding core ribosomal proteins such as *Rps21* and *Rpl37a*, suggesting high protein synthesis activity.

We also searched for expression of *Tgm2, Sema3f and Cxcl12*, which were recently found to mark three functional different epicardial populations in the developing zebrafish heart (3). We found *Tgm2* and *Sema3f* preferentially expressed in *Wt1*-expressing EpiSC-1 and 7 (Figure 2—figure supplement 2A-B). However, *Mylk*, which is an additional marker of the *Sema3f*-expressing zebrafish epicardial population (3), was primarily expressed in EpiSC-8. *Cxcl12*, present only in a small cell population in the zebrafish (3), was rather broadly expressed with enrichment in EpiSC-2 and subcluster EpiSC-1.1 (Figure 2—figure supplement 2C-D). These findings demonstrate some degree of evolutionary preservation in epicardial populations from zebrafish to mice, but also reveal considerable differences in marker gene expression pattern and population sizes.

### Spatial RNA expression of EpiSC population identifiers

To explore the specific location of EpiSC populations in the infarcted heart, RNA *in situ* hybridization of gene transcripts for selected EpiSC populations was carried out (Figure 2B upper panel), using population-specific identifiers (Figure 2B lower panel). *Msln* expression (EpiSC-7) was detected on the outer part of the epicardium, consistent with the epithelial signature of EpiSC-7. Expression of *Wt1* (EpiSC-7, 1, 8) and *Cd44* (EpiSC-4) also labelled cells that were localized on the outer layer. In contrast, *Sfrp2* expression (EpiSC-6) was rather homogenously dispersed throughout the epicardium and was also detected in the myocardium, labelling aCSC. A similar distribution pattern was observed for *Pcsk6* (EpiSC-3), *Top2a* (EpiSC-9; subpopulations of EpiSC-1, EpiSC-2) and *Dkk3* (EpiSC-10). In summary, we confirmed that major populations identified by scRNAseq localize to the activated post-MI epicardium.

### Cardiomyogenesis, paracrine factors and HIF-1-responsive genes

Lineage tracing experiments have shown that cardiomyogenesis can be initiated *in vivo* in Wt1^+^ epicardial cells when stimulated with thymosin β4 (10). We found thymosin β4 (*Tmsb4x*) and BRG1 (*Smarca4*), a transcription activator for WT1 (24), to be co-expressed within the *Wt1*-expressing EpiSC-8 (Figure 2C). Interestingly, many key cardiogenic factors were also expressed within the *Wt1*-expressing EpiSC populations (Figure 2C, cardiogenic factors I). This includes MESP1 which marks early cardiovascular progenitor specification (25), as well as WNT11, ISL1, TBX5 and GATA4, 5, and 6 which all play a critical role in heart development (26,27). Surprisingly, several *Hox* family members (Figure 2C) were expressed in EpiSC-8. HOX transcription factors are downstream effectors of retinoic acid signaling (28) that is required for differentiation of cardiac progenitors during heart development (29). Intriguingly, a second set of cardiogenic factors was found in EpiSC-3, 5, 6, 9 and 10 (Figure 2C, cardiogenic factors II). This includes Nkx-2.5 as well as BMP2 and BMP4 the latter of which are crucial in the regulation of Nkx-2.5 expression and specification of the cardiac lineage (30). In addition, we found low expression of genes encoding muscle structural proteins such as *Myl2, 4*, and *6, Tnnt2, Ttn* and *Nebl* which appear to be present especially in EpiSC-7 and EpiSC-9 (Figure 2C, contractile proteins). The highest expression levels of Notch target genes were found in *Wt1*-negative EpiSC populations, especially EpiSC-10 (Figure 2C). This is remarkable, since Notch-activated epicardial-derived WT1^+^ cells were described as a multipotent cell population with the ability to express cardiac genes (31). The observation that EpiSC-7 and 9 express cardiac specification markers and sarcomere proteins is suggestive that these populations have cardiomyogenic potential.

Epicardial cells have been reported to secrete numerous paracrine factors that can modulate myocardial injury in the mouse heart (8). In line with this observation, we found several chemokines (MCP-1, *Ccl2*; MIP-1β, *Ccl4*; RANTES, *Ccl5*; MCP-2, *Ccl8*) known to be involved in monocyte recruitment to be primarily expressed in EpiSC-11 (Figure 2D). EpiSC-11 also showed strongly enriched expression of IP-10 (*Cxcl10*, top 5 marker gene) which is involved in triggering anti-fibrotic effects after MI (32). Expression of chemokines involved in neutrophil granulocyte recruitment (GRO-α, β, γ, *Cxcl1, 2, 3*; ENA-78, *Cxcl5*; NAP-2/CXCL-7, *Ppbp*) was preferentially found in EpiSC-2 and 4 but also in EpiSC-1 and 7. Expression of SDF-1 (*Cxcl12*), known to reduce scar size when administered to the damaged heart (33), was enriched in EpiSC-2.

Expression of proangiogenic factors previously identified in the supernatant of WT1^+^ epicardial cells (8), such as Angiopoietin-1 (*Angpt1*), FGF2, IL-6, VEGFA and VEGFC, was not limited to the *Wt1*-expressing EpiSC populations but pertained to other identified EpiSC populations (Figure 2E). The most highly expressed paracrine factors were TGF-β1 which attenuates myocardial ischemia-reperfusion injury (34), IGF-1 which prevents long-term left ventricular remodeling after cardiac injury (35) and MYDGF which mediates ischemic tissue repair (36). A hypoxia-responsive element in the *Wt1* promotor was reported to bind HIF-1α that is required for WT1 induction (37). We found expression of *Hif1a* and HIF-1-responsive genes, particularly those encoding glycolytic enzymes, to be enriched in multiple EpiSC populations (Figure 2F).

### Post-MI aCSC in comparison to EpiSC

Clustering analysis of aCSC revealed 11 transcriptionally different populations (Figure3A, Figure 3—figure supplement 1A-C). Again, cell doublets with hybrid transcriptomes identified by DoubletFinder (16) (Figure 3—figure supplement 2A) and minor non-stromal cell populations (Figure 3—figure supplement 2B) were excluded from further analysis. The top 5 most differentially expressed genes in aCSC are displayed Figure 3—figure supplement 1D. Genes encoding ECM proteins were associated with aCSC-1 (*Eln, Wisp2*), aCSC-2 (*Thbs4, Cilp*), aCSC-4 (*Smoc2, Sparcl1*). Cell populations with similar gene signature were recently termed activated fibroblasts (11) or late response fibroblasts and matrifibrocytes (12). In these studies, a population similar to aCSC-5 was either referred to as Sca1-low or homeostatic epicardial-derived fibroblasts (11,12). aCSC-6 preferentially expressed genes characteristic of epithelial cells (*Lgals7, Dmkn, Msln*). Because this epithelial signature was comparable to that observed for EpiSC-7, it is very likely that cells of aCSC-6 are of epicardial origin and are due to incomplete removal during the cell isolation procedure. Consistent with this, *in situ* hybridization identified *Msln* expression exclusively within the epicardium (Figure 2B). Similar to so called cycling fibroblasts/proliferating myofibroblasts (11,12), we found genes involved in cell cycle and mitosis preferentially in aCSC-7 (*Cenpa, Stmn1, Ccnb2*) and aCSC-10 (*Top2a, Ube2c, Hist1h2ap, Pclaf*). Again similar to data in the literature (11,12), we found genes related to the cellular response to interferon highly expressed in aCSC-8 (*Ifit3, Isg15, Ifit1, Iigp1, Ifit3b*), which were termed interferon-stimulated/interferon-responsive fibroblasts. aCSC-9 highly expressed *Ly6a*, encoding Sca1, and resembled the reported Sca1-high/progenitor-like fibroblasts (11,12). Genes encoding contractile proteins were preferentially expressed in aCSC-3 (*Acta2, Tpm2*), which were recently referred to as myofibroblasts (12). Expression of Wnt pathway-associated genes was enriched in aCSC-1 (*Sfrp1, Sfrp2*) and aCSC-11 (*Wif1, Dkk3*), the latter of which were referred to as Wnt-expressing/endocardial-derived fibroblasts (11,12). Taken together, all aCSC populations identified by us are consistent with previously identified cardiac fibroblast populations.

**Figure 3.**
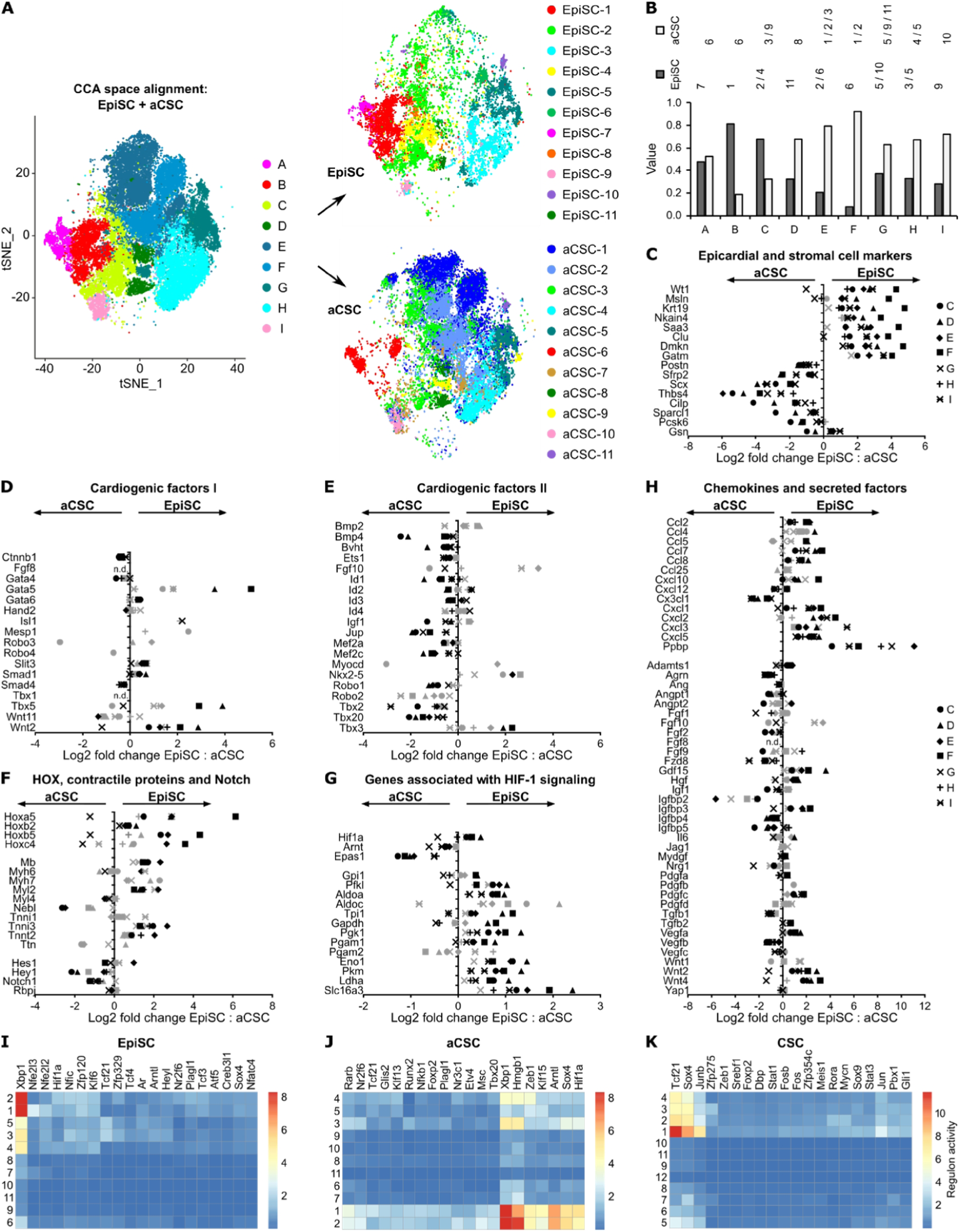
Comparison of EpiSC to aCSC. (**A**) Canonical correlation analysis (CCA) space alignment of EpiSC and aCSC scRNAseq data in one *t*-SNE plot (left) and split in one plot each (right). Cells are color-coded according to their assignment to CCA clusters (left) or previously identified populations (right). (**B**) Contribution of EpiSC and aCSC fractions to CCA clusters. (**C** to **H**) Relative expression of epicardial and stromal cell markers (C), cardiogenic factors (D and E), HOX transcriptions factors, contractile proteins and Notch target genes (F), genes associated with HIF-1 signaling (G) as well as chemokines and other secreted factors (H) in aCSC and EpiSC fractions as log2 fold change. Black symbols p-value ≤ 0.001; grey symbols p-value > 0.001. n.d., not defined. (**I** to **K**) Gene regulatory network analysis in EpiSC (I), aCSC (J) and CSC (K) populations by SCENIC.

To directly compare the transcriptional profile of EpiSC to myocardial aCSC, we performed canonical correlation analysis (CCA) space alignment of the two scRNAseq data sets (average RNA expression levels and differentially expressed marker genes are listed in Supplementary files 5 and 6, respectively). Generated CCA clusters (Figure 3A, left panel) are also displayed according to original cell IDs (Figure 3A, right panels). The contribution of EpiSC and aCSC fractions to CCA clusters is summarized in Figure 3B. As can be seen, CCA clusters B and C were mainly composed of EpiSC indicating that the transcriptional profile of EpiSC-1, 2, 4 is prevalent in the epicardium (Figure 3B). On the other hand, CCA clusters D-I were dominated by aCSC. Individual assignments showed that *Wt1*-negative EpiSC-3, 5, 6, 9, 10 carried to a variable degree expression signatures of genuine aCSC-1, 2, 3, 4, 5, 10, 11. Despite these similarities, there were multiple significant differences in average gene expression levels when comparing *Wt1*-negative EpiSC and aCSC within individual CCA clusters (Supplementary file 7). *Wt1*-expressing CCA clusters A and B were excluded from this comparative analysis, since they contained aCSC6, which probably represents epicardial contamination in the aCSC fraction (see above). As shown in Figure 3C, expression of epithelial/epicardial genes was generally higher in EpiSC, while mesenchymal/fibroblast genes were more dominant in aCSC. A distinct fraction of cardiogenic factors set I (see Figure 2C) was preferentially expressed in EpiSC (Figure 3D) while the opposite was true for cardiogenic factors set II (Figure 3E). Notably, cardiac contractile proteins (Figure 3F) and HIF-1-responsive glycolytic enzymes (Figure 3G) were predominantly expressed within EpiSC. Among the paracrine factors (Figure 3H) we found multiple chemokines highly enriched in EpiSC.

To further compare cell states in the two stromal cell fractions, we used SCENIC (18) for gene regulatory network reconstruction. This tool scores the activity of transcription factors by correlating their expression with the expression of their direct-binding target genes (18). As shown in Figure 3 I and J, EpiSC and aCSC showed distinct patterns of network activity, again emphasizing their different cellular identity and function.

Clustering analysis of CSC from uninjured hearts revealed 12 transcriptionally different populations (Figure 3—figure supplement 3A-C) after cell doublets (Figure 3—figure supplement 4A) and minor non-stromal cell populations (Figure 3—figure supplement 4B) were excluded. Interestingly, the smallest CSC population (CSC-12) highly expressed epithelial/epicardial genes (*Msln, Upk3b, Nkain4, Krt19*) (Figure 3—figure supplement 3D) and therefore is likely to represent cells of the epicardial monolayer. Individual single-cell analysis of this monolayer is technically not feasible, because the number of cells liberated by shear force from uninjured hearts is only rather small (estimated ∼ 5,600 cells/heart) and there is shear-independent release of cells (background) of unknown origin (13). As shown in Figure 3—figure supplement 5A-B, CCA space alignment of the CSC and EpiSC data set revealed that cells with the profile of EpiSC-1, 7 (CCA cluster a) and EpiSC-2, 4, 8 (CCA cluster c) were clearly prevalent in EpiSC, while the majority of CSC correlated with EpiSC-3, 5 and 6 (CCA cluster f, g, h). Direct comparison of EpiSC and CSC within individual CCA clusters again showed that expression of epithelial/epicardial genes was enriched in EpiSC, while transcript levels of the conventional fibroblast marker *Gsn* were higher in CSC (Figure 3—figure supplement 5C). As to be expected, gene regulatory network analysis by SCENIC revealed major differences of CSC to EpiSC and aCSC (Figure 3 I to K).

### Hierarchy of EpiSC populations

To better define the cellular relationship between the identified EpiSC populations, we combined scRNAseq of EpiSC with lineage tracing using tamoxifen-inducible Wt1-targeted reporter (Wt1^CreERT2^Rosa^tdTomato^) mice (Figure 4A). Transcriptional profiles of 13,373 cells from 2 mouse hearts 5 days after MI were assigned to previously defined EpiSC populations (Figure 4B). Surprisingly, expression of *tdTomato* fully overlapped with that of *Wt1* (Figure 4C), suggesting that WT1^+^ cells within the given time did not convert in cells of any *Wt1*-negative EpiSC population.

**Figure 4.**
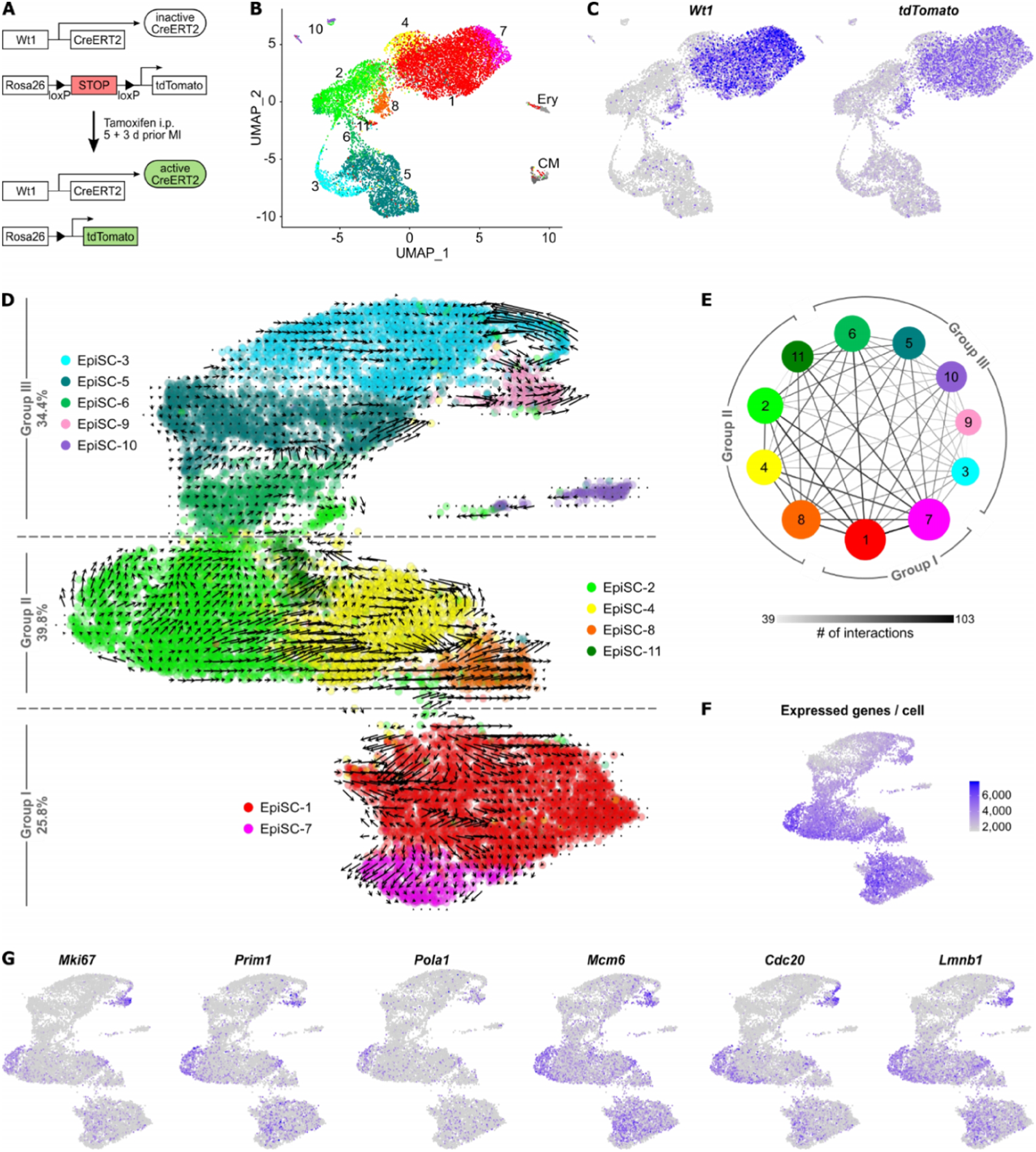
Cell hierarchy of EpiSC populations. (**A** to **C**) Lineage tracing of *Wt1*-expressing cell populations post MI using Wt1^CreERT2^Rosa^tdTomato^ mice. The experimental design is outlined in (A). UMAP plot of clustered scRNAseq data of the EpiSC fraction (*n*=13,373 single cells) pooled from two hearts 5 days post MI is shown in (B). Expression of *Wt1* and *tdTomato* is visualized in (C). (**D**) RNA velocity of EpiSC populations projected on the UMAP plot. Arrows show the local average velocity evaluated on a regular grid, indicating estimated future states. (**E**) Networks visualizing number of potential specific interactions between EpiSC populations as determined by CellPhoneDB. (**F**) Feature plot displaying the number of unique genes detected in each cell. (**G**) Expression of genes associated with cell proliferation visualized in feature plots.

RNA velocity analysis (19), which can predict and visualize future cell states based on the ratio of unspliced and spliced mRNA read counts, supports the notion that EpiSC consists of different independent groups of cell populations. As shown in Figure 4D, EpiSC-7 and 1 (group I) appeared to be separated from EpiSC-2, 4, 8 (group II) and EpiSC-6, 5, 3, 9 (group III).

To study cell-cell communication mediated by ligand-receptor interactions between the EpiSC populations, we used CellPhoneDB (20). As shown in Figure 4E, the highest numbers of interactions were predicted between groups I and II. Ligand-receptor pairs potentially involved in the interaction between the groups include WNT4, TGF-β2 and IGF1 signaling (Supplementary file 8). Group III not only showed the lowest cell-cell connections, it also showed the lowest number of expressed genes (Figure 4F). Since the expressed genes per cell correlate with developmental potential (38), this may indicate that a large fraction of group III cells are terminally differentiated.

Mapping of genes characteristic for active cell cycle progression (39) in feature plots of EpiSC showed that each of the three EpiSC groups identified above was equipped with a cell cluster expressing a number of cell cycle genes (Figure 4G). Within group I it is subcluster EpiSC-1.4, within group II a subpopulation of EpiSC-2 and within group III EpiSC-9. This location is very similar to the origin of velocity arrows (Figure 4D).

## Discussion

In this study we provide a single-cell landscape of the post-MI epicardium with 11 transcriptionally different cell populations. This amazing degree of cellular heterogeneity is similar to that of myocardial cardiac fibroblasts (11,12). The widely used epicardial lineage marker gene *Wt1* (4) was selectively expressed in three populations (EpiSC-1, 7, 8), which were localized on the outer surface of the activated epicardium (Figure 2B). Other commonly used lineage markers and recently identified epicardial population markers of the developing zebrafish heart (3) were rather heterogeneously distributed (Figure 2A, Figure 2—figure supplement 2A-B), suggesting that they mark different stages of differentiation, commitment or activity in the adult heart.

The quality of scRNAseq critically depends on the cell isolation technique which ideally should preserve the native state of cells as close as possible. To minimally perturb the native expression profile of cardiac stromal cells we have recently elaborated a perfusion protocol for simultaneous isolation of EpiSC and aCSC from the same infarcted heart, which is short (8 min) and results in a high yield of viable cells (13). The application of mild shear forces on the cardiac surface by a simple motor-driven device permitted the rather selective removal of the EpiSC fraction (13). As compared to a commonly used mincing protocol (30-40 min), we found the yield of aCSC to be considerably higher with our technique and this was associated with significant lower induction of immediate early response genes (13). Because of these reasons our data are difficult to compare with published studies at single-cell resolution of the post-MI heart which relied on a mincing protocol. These studies have identified either no (40–42) or only one (11,12,43,44) epicardial cell population.

That there are different and genuine cellular identities of EpiSC and aCSC is supported by CCA space alignment analysis and gene regulatory network analysis, which revealed distinct expression of epicardial *vs*. mesenchymal/fibroblast genes (Figure 3C) as well as various paracrine factors (Figure 3H) and individual patterns in transcription factor activity (Figure 3 I and J), respectively. Direct comparison of cellular expression signatures between EpiSC and aCSC also revealed many transcriptional similarities which is not surprising in view of their close developmental relationship (45).

Cellular distribution of cell cycle genes (Figure 4G) and RNA velocity analysis (Figure 4D) suggest that the EpiSC populations can be classified into three different population groups, each containing a cluster of proliferating cells. Marker genes of group I, comprising *Wt1*-expressing EpiSC-1 and 7, have been reported in healthy adult mouse epicardium obtained by laser capture (23) and have been exclusively used in previous single-cell studies to identify epicardial cells (27, 28) (Figure 1F). Surprisingly, group I cells accounted for only 26% of the EpiSC fraction from the injured heart. Furthermore, the expression of several paracrine proangiogenic factors that were previously considered specifically derived from WT1^+^ epicardial cells (8) was not limited to group I EpiSC, but was even higher in other EpiSC populations (Figure 2E). Thus, about 2/3 of all epicardial cells were *Wt1*-negative, but they are likely to be also involved in the secretion of paracrine factors. Furthermore, aCSC expressed numerous paracrine factors (Figure 3H). Therefore, secretion of paracrine factors appears to be general feature of both epicardial and myocardial stromal cells which all can contribute to cardioprotection.

Group II, consisting of four cell populations, represented 40% of the EpiSC fraction and showed enriched expression of chemokines known to be involved in attraction of monocytes and neutrophils (Figure 2D). The expression of these chemokines was a general feature of EpiSC in comparison to aCSC (Figure 3H). This finding points to a role of epicardial cells in the modulation of the innate immune response post MI, as was already suggested with regard to adaptive immune regulation during the post-MI recovery phase (9).

Cells with the transcriptional profiles of group I and II populations were generally prevalent in EpiSC in comparison to aCSC (Figure 3B). Group I and II populations also showed the highest number of potential ligand-receptor interactions (Figure 4E). Interestingly, EpiSC-8, characterized by expression of *Wt1* and high transcript levels of HOX transcription factors as well as of genes annotated with the GO term “Embryonic organ morphogenesis”, also specifically expressed thymosin β4 (Figure 2C) which has tissue-regenerating properties (10). The pronounced expression of cardiogenic factors and contractile proteins in group I (Figure 2C), and the parallel expression of WT1 and thymosin β4 suggests that this cellular network has cardiogenic potential and was involved in the previously reported formation of cardiomyocytes from WT1^+^ cells after thymosin β4 stimulation (10). Interestingly, group I and II also shared high expression of HIF-1-responsive glycolytic enzymes (Figure 2F), which again was a general feature of EpiSC compared to aCSC (Figure 3G). This is in support of the epicardium being a hypoxic niche (46). Since WT1 expression is HIF-1-dependent (37), this is consistent with the view that epicardial HIF-1 signaling is likely to be an important trigger in the ischemic heart to promote cardioprotection.

Group III comprised five cell populations accounting for 34% of all EpiSC. In contrast to group I, group III EpiSC were found localized throughout the activated epicardium (Figure 2B) and group III population identifiers also labeled stromal cells within the myocardium (Figure 2B). This finding is consistent with the transcriptional profile of group III EpiSC that was quite similar to that of major aCSC populations (Figure 3A and B) and the CSC fraction (Figure 3—figure supplement 5 A-B). This demonstrated that group III cells exhibit a fibroblast-like phenotype. In addition, group III EpiSC showed the lowest numbers of both predicted cell-cell interactions (Figure 4E) and expressed a lower number of genes (Figure 4F), suggesting a less active cell state. Another remarkable feature of group III was the expression of a second set of cardiogenic factors (Figure 2C). However, these cardiogenic factors were prevalently expressed in aCSC (Figure 3E) which is consistent with the postulated cardiogenic potential of cardiac fibroblasts (47). As to the cellular origin of EpiSC, our lineage tracing data suggest, that at day 5 post MI WT1^+^ cells are not the major contributor to epicardial expansion. The *Wt1*-negative fibroblast-like cells in group III may have either been derived from group II cells or they are myocardial stromal cells that have migrated into the epicardial multi-cell layer.

Histological observations by Zhou *et al*. (8) and Quijada *et al*. (4) found no evidence for migration of WT1^+^ cells into the infarcted myocardium even after longer periods of time. That WT1^+^ cells most likely do not migrate into the infarcted heart appears to be in contrast with a study using lentiviral labeling of epicardial cells after pericardial injection of virus expressing fluorescent protein (48). Along the same line we have previously shown, that tracking of epicardial cells after labeling with fluorescently marked nanoparticles revealed migration into the injured heart (49). Since group II and III EpiSC constitute the majority of post-MI epicardial cells, it is well conceivable that populations of *Wt1*-negative cells were preferentially marked by the above-mentioned labeling techniques and this may explain the reported migration into the injured myocardium.

In summary, our study explored post-MI epicardial cell heterogeneity in the context of all cardiac stromal cells at unprecedented cellular resolution. Important epicardial properties in post-MI wound healing/regeneration can now be attributed to specific cell populations. A deeper understanding of adult epicardial hierarchy may help to decipher the signaling mechanisms by which individual epicardial cell populations interact to specifically stimulate cardiac repair processes originating in the epicardium.

## Methods

### Mice

All animal experiments were performed in accordance with the institutional guidelines on animal care and approved by the Animal Experimental Committee of the local government “Landesamt für Natur, Umwelt und Verbraucherschutz Nordrhein-Westfalen” (reference number 81-02.04.2019.A181). The animal procedures conformed to the guidelines from Directive 2010/63/EU of the European Parliament on the protection of animals used for scientific purposes. For this study, male C57BL/6 (Janvier, Le Genest-Saint-Isle, France) and male tamoxifen-inducible Wt1-targeted (Wt1^CreERT2^Rosa^tdTomato^) mice were used. Inducible WT1^CreERT2^Rosa^tdTomato^ were generated by crossing the Wt1^tm2(cre/ERT2)Wtp^/J strain (stock no: 010912 Jackson Laboratory, Bar Harbor, US) with the B6;129S6-Gt(ROSA)26Sor^tm14(CAG-tdTomato)Hze^/J strain (stock no. 007908; Jackson Laboratory, Bar Harbor, US) followed by genotyping. Mice (body weight, 20-25 g; age, 8-12 weeks) used in this study were housed at the central animal facility of the Heinrich-Heine-Universität Düsseldorf (ZETT, Düsseldorf, Germany), were fed with a standard chow diet and received tap water *ad libitum*.

### Animal procedures

Myocardial infarction (MI) followed by reperfusion was performed as previously described (14). In brief, mice were anaesthetized (isoflurane 1.5%) and the left anterior descending coronary artery (LAD) was ligated for 50 min followed by reperfusion. LAD occlusion was controlled by ST-segment elevation in electrocardiography recordings. Sham control animals underwent the surgical procedure without LAD ligation.

Wt1^CreERT2^-mediated lineage tracing was performed as described previously (10). In brief, Wt1^CreERT2^Rosa^tdTomato^ mice received tamoxifen injections (2 mg emulsified in sesame oil; i.p.) 5 and 3 days prior MI to induce CreERT2 activity.

### Isolation of EpiSC, aCSC and CSC

EpiSC and aCSC at day 5 after MI and control CSC from uninjured hearts at day 5 after sham surgery were isolated as previously described (13). In brief, mice were sacrificed by cervical dislocation and hearts were excised for preparation of the aortic trunk in ice-cold PBS. Isolated hearts were immediately cannulated and perfused with PBS (3 min), followed by perfusion with collagenase solution (8 min; 1,200 U/ml collagenase CLS II (Biochrom, Berlin, Germany) in PBS at 37°C.

EpiSC were simultaneously isolated from post-MI hearts by bathing the heart in its collagenase-containing coronary effluat while applying mild shear force to the cardiac surface. The effluat was collected, centrifuged (300 g, 7 min) and cells were resuspended in PBS / 2% FCS / 1 mM EDTA. Application of shear force and effluat collection was omitted for uninjured sham control hearts due to the absence of an activated, expanded epicardial layer.

aCSC and CSC were isolated by mechanical dissociation of the digested myocardial tissue, followed by resuspension in Dulbecco’s modified eagle medium (DMEM) / 10% FCS. The cell suspensions were meshed through a 100 μm cell strainer and centrifuged at 55 g to separate cardiomyocytes from non-cardiomyocytes. The supernatants were again passed through a 40 µm cell strainer, centrifuged (7 min, 300 g) and cell pellets were resuspended in PBS / 2% FCS / 1 mM EDTA. Cells were immediately stained for surface markers and applied to fluorescence-activated cell sorting.

### Fluorescence-activated cell sorting

Cell fractions (EpiSC, aCSC, CSC) were isolated as described above and stained at 4°C (15 min) with fluorochrome-conjugated antibodies against surface markers of endothelial cells (CD31 (BD Biosciences, Franklin Lakes, USA) and myeloid cells (CD45 (BD Biosciences)) in presence of 7AAD (viability marker (BD Biosciences)). Sorting was performed with a MoFlo XDP flow cytometer (Beckman Coulter, Brea, USA), where dead cells (7AAD^+^) and CD31^+^/CD45^+^ cells were excluded by gating on 7AAD^-^/CD31^-^/CD45^-^cells. Sorting of cells isolated from the Wt1^CreERT2^Rosa^tdTomato^ mouse line followed the same gating strategy with one minor change. Fixable Viability Dye eFluor 780 (eBioscience, San Diego, USA) was used instead of 7AAD to avoid fluorescence spill-over.

### ScRNAseq

The sorted single-cell suspensions were directly used for the scRNAseq experiments. ScRNAseq analysis was performed by using the 10x Genomics Chromium System (10x Genomics Inc San Francisco, CA). Cell viability and cell number analysis were performed via trypan blue staining in a Neubauer counting chamber. A total of 2,000 to 20,000 cells, depending on cell availability, were used as input for the single-cell droplet libraries generation on the 10x Chromium Controller system utilizing the Chromium Single Cell 3’ Reagent Kit v2 according to manufacturer’s instructions. tdTomato lineage tracing experiments were conducted utilizing the Chromium Single Cell 3’ Reagent Kit v3 according to manufacturer’s instructions. Sequencing was carried out on a HiSeq 3000 system (Illumina Inc. San Diego, USA) according to manufacturer’s instructions with a mean sequencing depth of ∼90,000 reads/cell for EpiSC and ∼70,000 reads/cell for aCSC. Differences in sequencing depth were necessary in order to achieve a similar sequencing saturation between all samples of ∼70%.

### Processing of scRNAseq data

Raw sequencing data was processed using the 10x Genomics CellRanger software (v3.0.2) provided by 10x Genomics. Raw BCL-files were demultiplexed and processed to Fastq-files using the CellRanger *mkfastq* pipeline. Alignment of reads to the mm10 genome and UMI counting was performed via the CellRanger *count* pipeline to generate a gene-barcode matrix.

The median of detected genes per cell was 3,155 for EpiSC, 3,265 for aCSC and 2,241 for CSC. The median of UMI counts per cell was 10,689 for EpiSC, 11,110 for aCSC and 5,879 for CSC. Mapping rates (reads mapped to the genome) were about 89% for EpiSC, 90.9% for aCSC and 86.7% for CSC.

For tdTomato lineage tracing experiments a custom reference, consisting of the mm10 genome and the full-length sequence of tdTomato, was generated via CellRanger *mkref*.

### Filtering and clustering of scRNAseq data

Further analyses were carried out with the Seurat v3.0 R package (15). Initial quality control consisted of removal of cells with fewer than 200 detected genes as well as removal of genes expressed in less than 3 cells. Furthermore, cells with a disproportionately high mapping rate to the mitochondrial genome (mitochondrial read percentages >5.0 for EpiSC and aCSC, >7.5 for CSC) have been removed, as they represent dead or damaged cells. Normalization has been carried out utilizing SCTransform. Biological replicates have been integrated into one dataset by identifying pairwise anchors between datasets and using the anchors to harmonize the datasets. Dimensional reduction of the data set was achieved by Principal Component analysis (PCA) based on identified variable genes and subsequent UMAP embedding. The number of meaningful Principal Components (PC) was selected by ranking them according to the percentage of variance explained by each PC, plotting them in an “Elbow Plot” and manually determining the number of PCs that represent the majority of variance in the data set. Cells were clustered using the graph-based clustering approach implemented in Seurat v3.0. Doublet identification was achieved by using the tool DoubletFinder (v2.0.2) (16) by the generation of artificial doublets, using the PC distance to find each cell’s proportion of artificial k nearest neighbors (pANN) and ranking them according to the expected number of doublets. Heat maps were generated using Morpheus (https://software.broadinstitute.org/morpheus).

### Gene Ontology enrichment analysis

Enriched Gene Ontology (GO) terms in differentially expressed genes between populations were identified by using the gene ontology enrichment analysis and visualization (GOrilla) tool (17).

### Canonical correlation analysis (CCA) space alignment

A direct comparison of the EpiSC and aCSC as well as EpiSC and CSC data sets was performed by Seurat’s CCA alignment procedure (v 2.3.4). Briefly, the top 600 variable genes were identified for each data set and subjected to a CCA. Herein, canonical correlation vectors were identified and aligned across data sets with dynamic time warping. After alignment, a single integrated clustering was performed, allowing for comparative analysis of cell populations across both cell fractions.

### Gene regulatory network analysis

Gene regulatory network reconstruction and cell-state identification in EpiSC, aCSC and CSC datasets was performed using the SCENIC (18).

### RNA velocity analysis

The Python software velocyto.py (Version 0.17) (19) was run on the EpiSC count matrices and BAM files generated by CellRanger (see above) to predict and visualize future cell states based on the ratio of unspliced and spliced mRNA read counts. For the visualization, cell clusters, PCA and UMAP data were imported by the Seurat analysis (see above).

### Cell-cell communication analysis

Cell-cell communication mediated by ligand-receptor complexes between EpiSC populations was analyzed using the tool CellPhoneDB v.2.0 (20) after mapping mouse genes to human orthologs.

### RNA *in situ* hybridization

*In situ* detection of selected marker gene expression was performed by RNA *in situ* hybridization using the RNAScope 2.5 HD Detection Kit Assay - RED (Advanced Cell Diagnostics, Hayward, CA) (21). Fresh frozen hearts (5 days after MI) were cut in 10 μm sections. Fixation and pretreatment of the cryosections were performed according to the manufacturer’s instructions. The incubation time for the hybridization of RNAScope 2.5 AMP 5 - RED was increased to 120 min to enhance the signal. For evaluation of the target probe signal the hybridized heart sections were examined under a standard bright field microscope (BX61 Olympus, Hamburg, Germany) using a 20x objective. Images were processed for publication using ImageJ/Fiji (22).

### Statistics

Markers defining each cluster as well as differential gene expression between different clusters were calculated using a two-sided Wilcoxon Rank-Sum test which is implemented in Seurat.

## Acknowledgements

We thank K. Raba (Institute of Transplantation Diagnostics and Cell Therapeutics, University Duesseldorf) for technical assistance with cell sorting by flow cytometry and J. Schulze (Department for Genomics & Immunoregulation, Limes, Bonn) for helpful advice and discussions. JS, JF, MG, NG were supported by a grant of the German Research Council (DFG, SFB1116, project identifier: 236177352). CO was supported by the DFG-funded International Research Training Group 1902 (project identifier: 220652768) and this work was part of his PhD thesis. JH was supported by the Research Committee of the Medical Faculty of the Heinrich Heine University Düsseldorf (project identifier: 2018-12).

## Author contributions

JH and CO conducted the experiments and wrote the manuscript. TL and KK conducted the single-cell measurements including sequencing and bioinformatic analysis using the 10x Genomics platform. RZ did the RNA *in situ* hybridization experiments. JW, AL, GK and CD performed bioinformatic analyses. ZD performed the animal handling and myocardial infarction. CA assisted with the experiments. MG, NG, JF and AL helped designing the experiments and discussed and/or interpreted findings. JS planned and coordinated the experiments, wrote and edited the manuscript.

## Data availability

ScRNAseq data have been deposited in the ArrayExpress database at EMBL-EBI (www.ebi.ac.uk/arrayexpress) under accession number E-MTAB-10035.

## Figure supplements titles/legends

**Figure 1—figure supplement 1.**
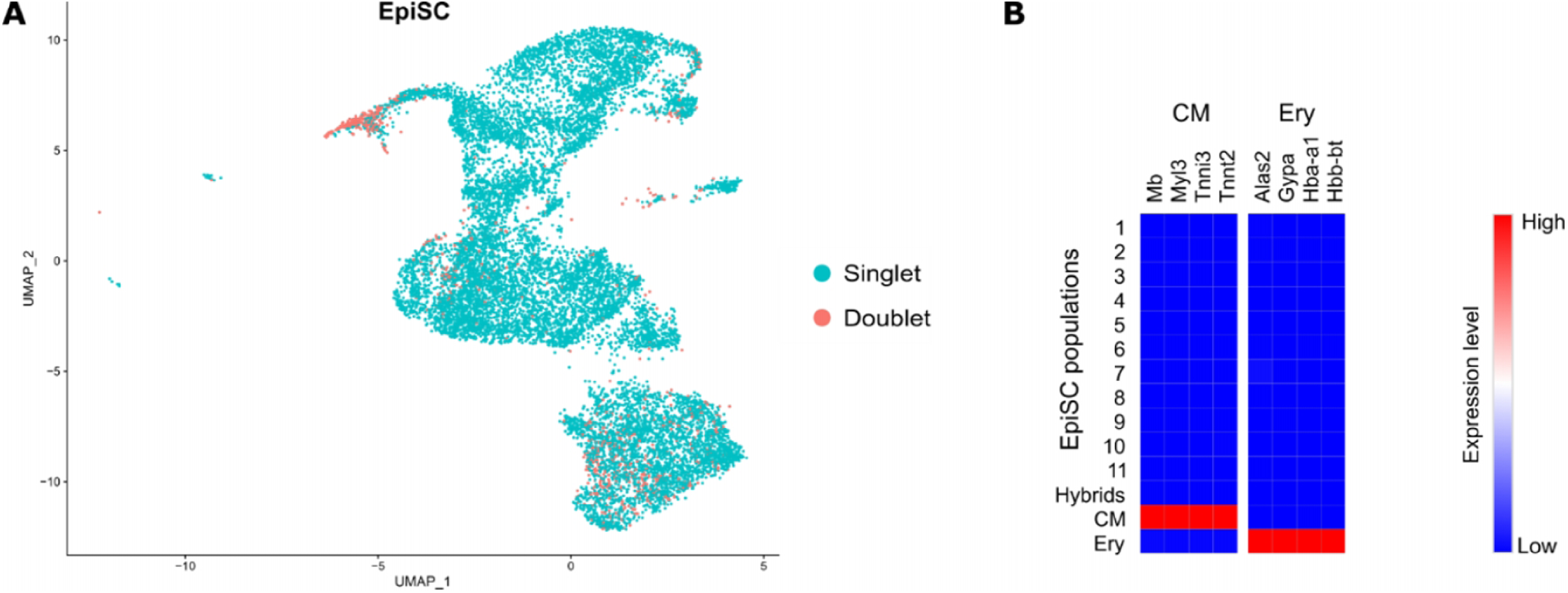
Excluded hybrid and non-stromal cell populations in the EpiSC fraction. (**A**) DoubletFinder tool was used to detect cell doublets with hybrid transcriptomes. **B** Heat map showing the expression of markers for cardiomyocytes (CM) and erythrocytes (Ery) used to identify residual populations of non-stromal cells.

**Figure 2—figure supplement 1.**
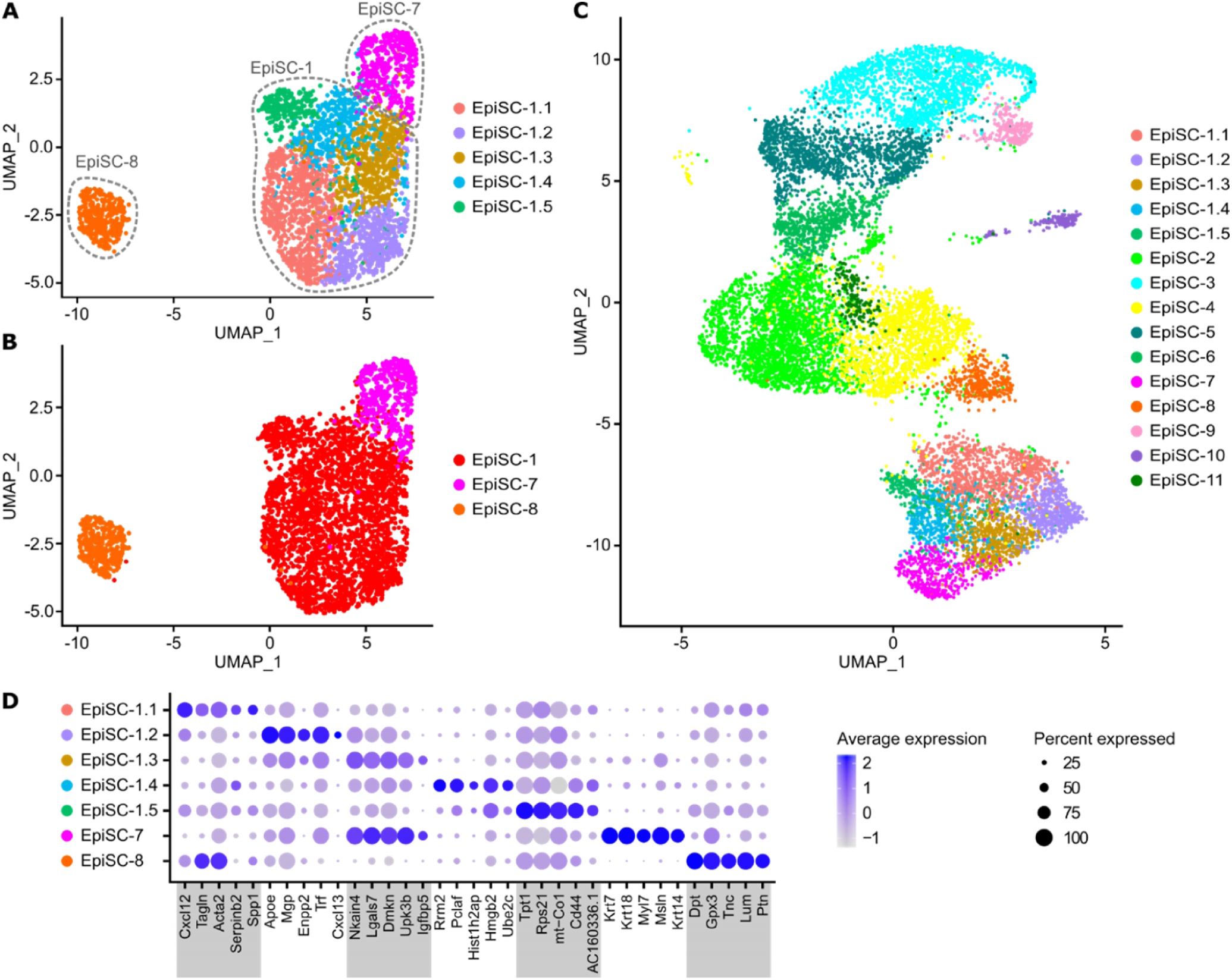
Subclusters of *Wt1*-expressing EpiSC populations. (**A**) UMAP plot of subclusters (Sub) of *Wt1*-expressing EpiSC-7, -1 and 8. (**B**) UMAP plot of the subclustering with labelling according to previous identity. (**C**) Transfer of subcluster labelling to UMAP plot of all EpiSC populations showing the position of identified subclusters. (**D**) Dot plot of top 5 marker genes for each subcluster together with EpiSC-7 and EpiSC-8.

**Figure 2—figure supplement 2.**
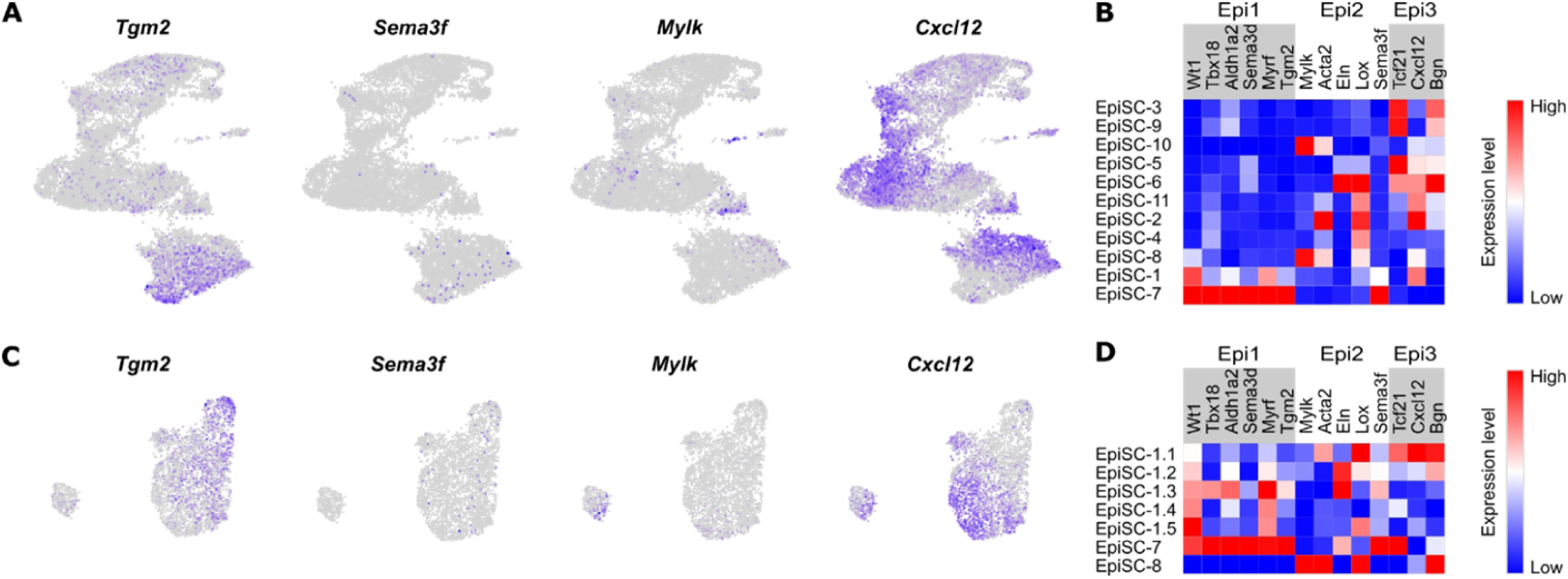
Expression of epicardial markers previously identified in the developing zebrafish heart in mouse post-MI EpiSC. Expression of epicardial population markers identified in the developing zebrafish heart by Weinberger *et al*. (3) visualized in the whole EpiSC fraction (**A** and **B**) and in *Wt1*-expressing subclusters (**C** and **D**) via feature plots and heat maps. For each of the three zebrafish epicardial populations (Epi1, Epi2, Epi3) selected marker genes are shown.

**Figure 3—figure supplement 1.**
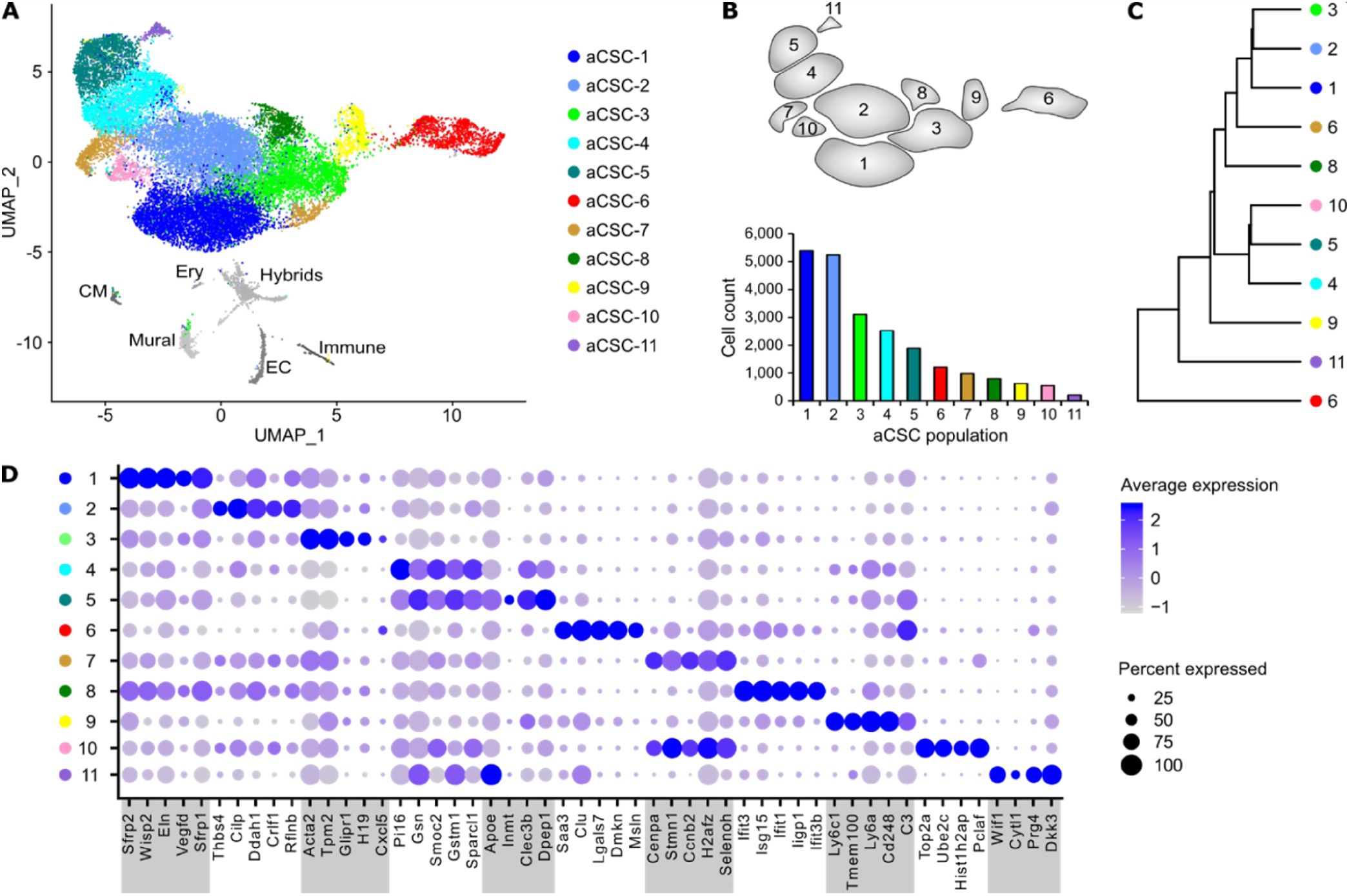
Cell populations in aCSC from infarcted myocardium. (**A**) UMAP plot of clustered scRNAseq data of the pooled aCSC fraction (*n*=24,470 single cells). Each point represents a single cell and identified cell populations are color-coded. CM, cardiomyocytes; Ery, erythrocytes; EC, endothelial cells. (**B**) Scheme of the populations from (A) and population cell counts of aCSC. (**C**) Dendrogram of aCSC populations according to average RNA expression. (**D**) Dot plot of top 5 marker genes for each aCSC population.

**Figure 3—figure supplement 2.**
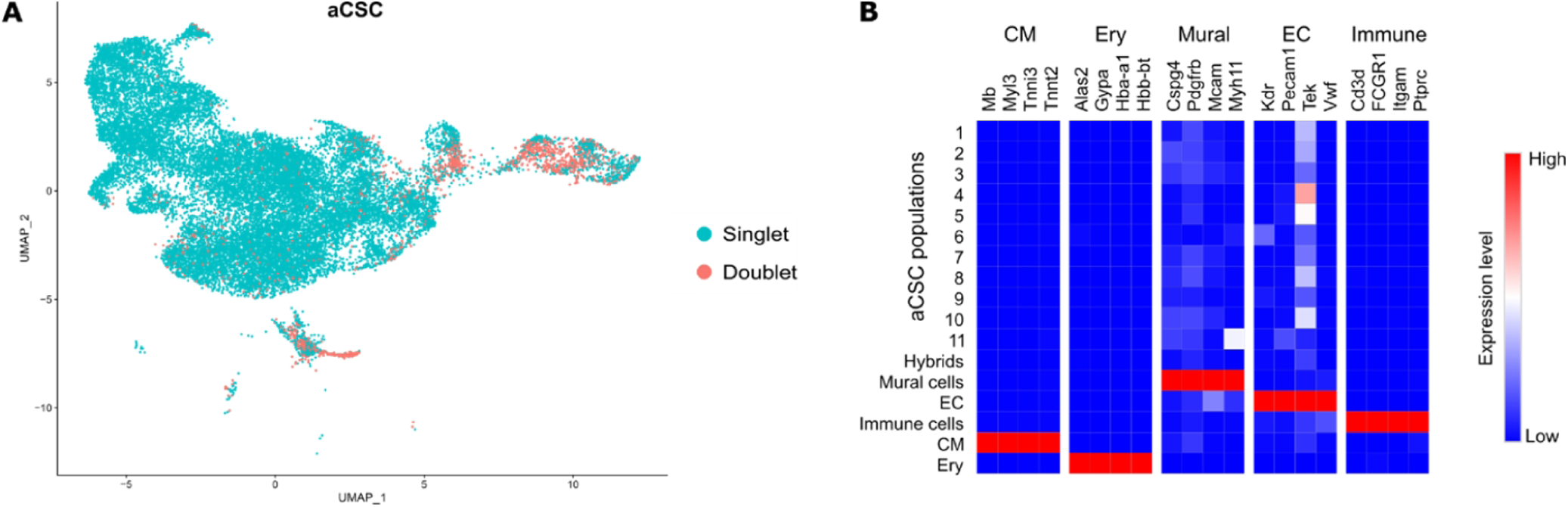
Excluded hybrid and non-stromal cell populations in the aCSC fraction. (**A**) DoubletFinder tool was used to detect cell doublets with hybrid transcriptomes. **B** Heat map showing the expression of markers for mural cells, endothelial cells (EC), immune cells, cardiomyocytes (CM), and erythrocytes (Ery) used to identify residual populations of non-stromal cells.

**Figure 3—figure supplement 3.**
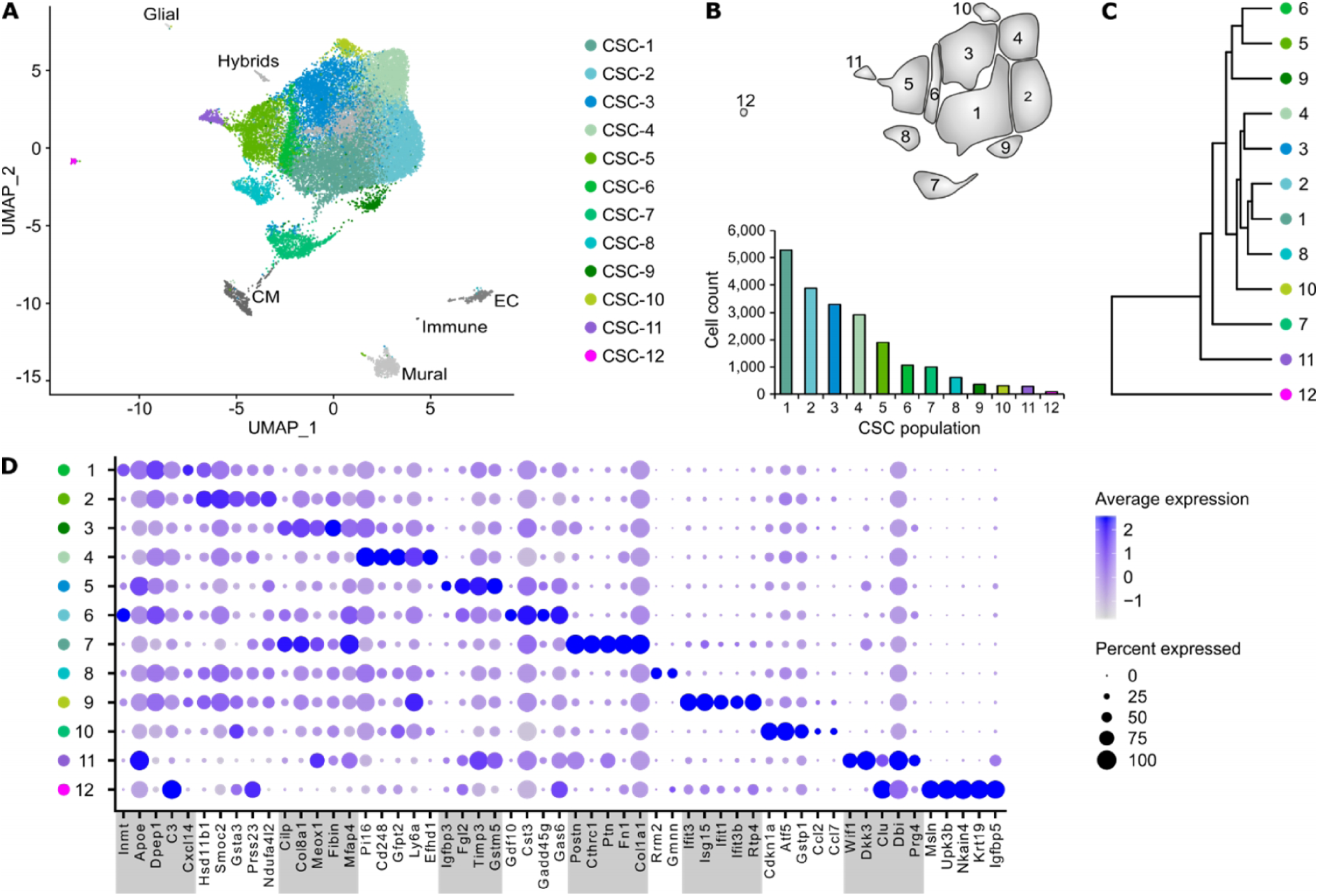
Cell populations in CSC from control hearts. (**A**) UMAP plot of clustered scRNAseq data of the pooled CSC fraction (*n*=24,781 single cells). Each point represents a single cell and identified cell populations are color-coded. CM, cardiomyocytes; EC, endothelial cells. (**B**) Scheme of the populations from (A) and population cell counts of and CSC. (**C**) Dendrogram of and CSC populations according to average RNA expression. (**D**) Dot plot of top 5 marker genes for each CSC population.

**Figure 3—figure supplement 4.**
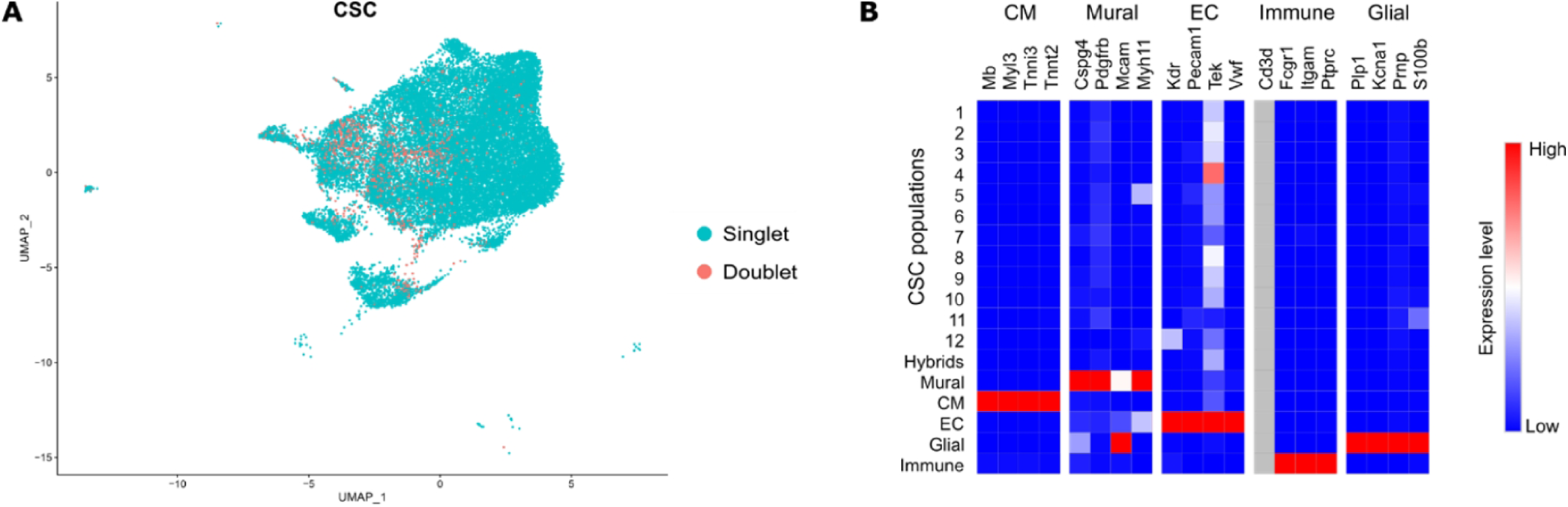
Excluded hybrid and non-stromal cell populations in the CSC fraction. (**A**) DoubletFinder tool was used to detect cell doublets with hybrid transcriptomes. (**B**) Heat map showing the expression of markers for mural cells, cardiomyocytes (CM), endothelial cells (EC), glial cells, and immune cells used to identify residual populations of non-stromal cells (grey filling, gene expression not detected).

**Figure 3—figure supplement 5.**
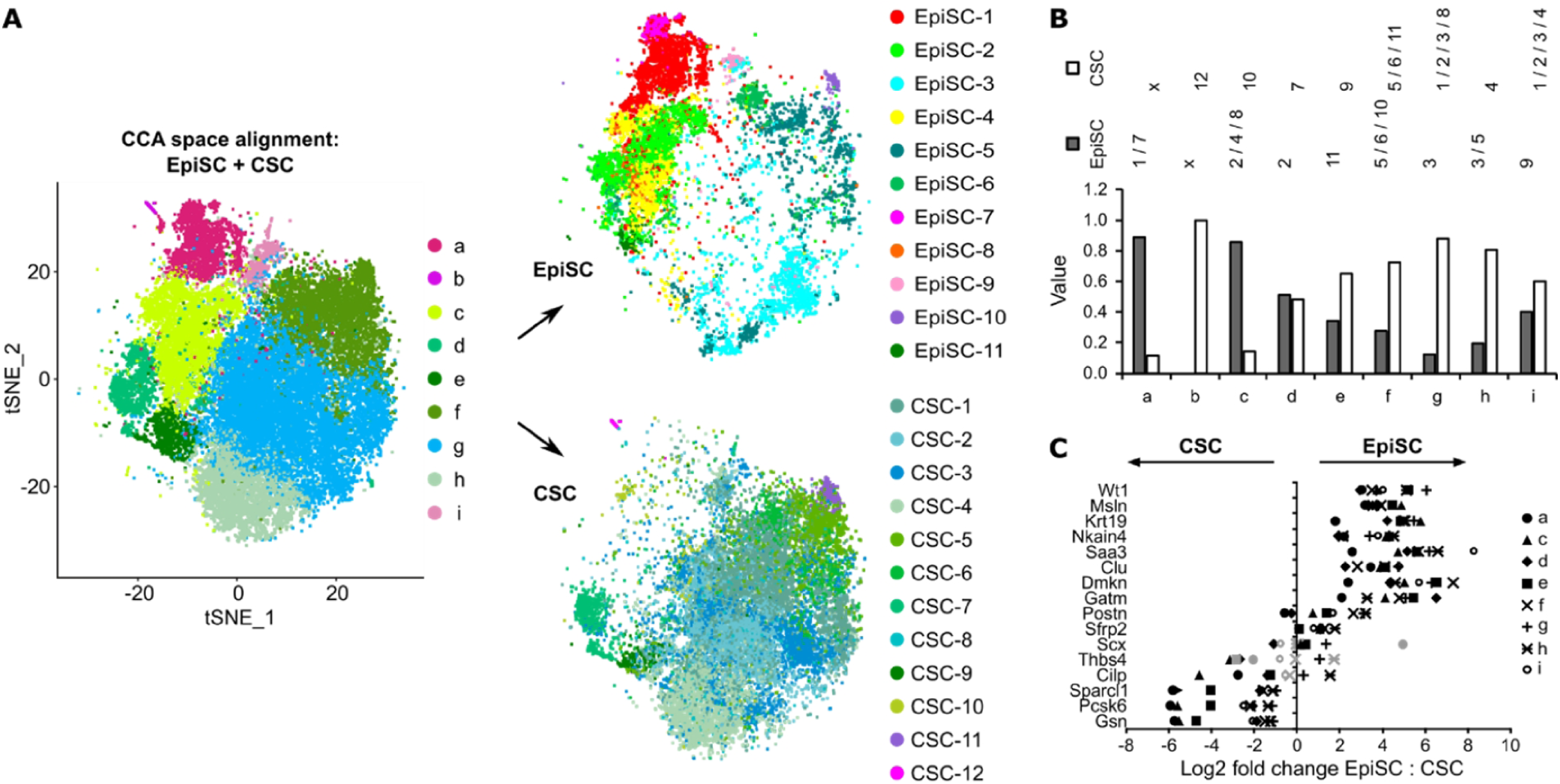
Comparison of EpiSC to CSC. (**A**) CCA space alignment of EpiSC and CSC scRNAseq data in one *t*-SNE plot (left) and split in one plot each (right). Cells are color-coded according to their assignment to CCA clusters (left) or previously identified populations (right). (**B**) Contribution of EpiSC and CSC fractions to CCA clusters. (**C**) Relative expression of epicardial and stromal cell markers in CSC and EpiSC fractions from non-epithelial CCA clusters C-I as log2 fold change. Black symbols p-value ≤ 0.001; grey symbols p-value > 0.001.

## Source data titles

**Figure 1**—**source data 1**

**Source data for EpiSC population cell counts summarized in Figure 1C**.

**Figure 3**—**source data 1**

**Source data for aCSC population cell counts summarized in Figure 3**—**Figure Supplement 1B**.

**Figure 3**—**source data 2**

**Source data for aCSC population cell counts summarized in Figure 3**—**Figure Supplement 3B**.

## Supplementary files titles

**Supplementary file 1**

**Average gene expression levels in EpiSC, aCSC and CSC populations**.

**Supplementary file 2**

**Genes with significantly enriched expression among EpiSC, aCSC and CSC populations**.

**Supplementary file 3**

**Average gene expression levels in subclusters of *Wt1*-expressing EpiSC populations**.

**Supplementary file 4**

**Genes with significantly enriched expression among subclusters of *Wt1*-expressing EpiSC populations. Supplementary file 5**

**Average gene expression levels in clusters from CCA space alignment of EpiSC and aCSC with separation in EpiSC and aCSC**.

**Supplementary file 6**

**Genes with significantly enriched expression among clusters from CCA space alignment of EpiSC and aCSC**.

**Supplementary file 7**

**Relative gene expression in EpiSC and aCSC within individual CCA clusters as log2 fold change with p-values**.

**Supplementary file 8**

**Selected ligand-receptor interactions between EpiSC populations as predicted by CellPhoneDB**.

